# Permanent neuroglial remodeling of the retina following infiltration of CSF1R-inhibition resistant peripheral monocytes

**DOI:** 10.1101/307900

**Authors:** Eleftherios I Paschalis, Fengyang Lei, Chengxin Zhou, Vassiliki Kapoulea, Reza Dana, James Chodosh, Demetrios G. Vavvas, Claes H. Dohlman

## Abstract

Previous studies have demonstrated that ocular injury can lead to prompt infiltration of bone marrow-derived peripheral monocytes into the retina. However, the ability of these cells to integrate into the tissue and become microglia has not been studied. Here we show that such peripheral monocytes not only infiltrate into the retina after ocular injury, but that they engraft permanently, migrate to the three distinct microglia strata, and adopt a microglia-like morphology. However, contrary to the original microglia, after injury the engrafted peripheral monocytes are resistant to depletion by colony stimulating factor 1 receptor (CSF1R) inhibitor and remain pro-inflammatory, expressing high levels of major histocompatibility complex II (MHC-II) for the long-term. In the absence of ocular injury, on the other hand, the peripheral monocytes that repopulate the retina after CSF1R inhibition remain sensitive to CSF1R inhibition and can be re-depleted. The observed permanent neuroglia remodeling after injury constitutes a major potential immunological change that may contribute to progressive retinal degeneration. These findings may be relevant also to other degenerative conditions of the retina and central nervous system.

**Significance statement:** Ocular injury causes permanent neuroglia remodeling that promotes neuroinflammation.

## Introduction

We recently showed that ocular injury causes prompt infiltration of peripheral monocyte into the retina (1). These cells gradually differentiate from ameboid c-c chemokine receptor 2^+^ (CCR2^+^) monocytes to highly ramified CX3C chemokine receptor 1^+^ (CX3CR1^+^) expressing macrophages (1) and acquire a morphology that resembles microglia. However, how these cells functionally integrate into the retina has not been investigated. Understanding the contribution of peripheral monocytes in retinal neuroglia remodeling may be clinically important in our understanding of mechanisms involved in progressive neuroretinal degeneration, especially after ocular injuries, since new depletion/antagonism strategies may be employed therapeutically for patients with neuroglia disorders (2, 3).

Peripheral monocytes/macrophages (Mφs) share common lineage with microglia which complicates the study of the subsequent differentiation of the two cell populations (4). Recently, new markers have been proposed for microglia labeling (5, 6); however, they still require further validation. Traditionally, studies have involved the use of transgenic mice and bone marrow chimeras, but such studies have often generated confusion, primarily due to side effects from gramma irradiation used in bone marrow transfer (BMT) which can result in blood-brain barrier disruption (4, 7-9) and peripheral monocyte infiltration. Thus, the presence of peripheral monocytes after bone marrow transfer was initially attributed to a physiological microglia turnover by peripheral monocytes. However, improved fate mapping techniques have now confirmed that peripheral monocytes/macrophages (Mφs) do not enter into the CNS or retina parenchyma under normal conditions (1, 10, 11); therefore, such infiltration happens only in pathologic conditions (8) and likely contributes to neuroinflammation (1, 12, 13).

This study was designed to examine the ontogeny and function of peripheral infiltrating macrophages into the retina after ocular injury (14). To address these questions we employed our previously characterized busulfan myelodepletion bone marrow chimera model for long-term fate mapping of peripheral monocyte infiltration into the retina (1). In addition, we used a small molecule inhibitor of colony stimulating factor 1 receptor (CSF1R) to perform a set of microglia depletion experiments to study the roles of microglia and peripheral monocytes in retinal repopulation. This study shows that peripheral monocytes are not only responsible for acute retinal inflammation after ocular injury (1, 14), but that they contribute to the permanent remodeling of the neuroglia system by establishing a pro-inflammatory phenotype that may promote or contribute to neurodegeneration. These findings may be clinically important and can help us better understand the mechanisms by which patients with acute ocular injuries (chemical, surgical, or traumas) become susceptible to progressive neurodegeneration processes, such as progressive retina ganglion cell loss (glaucoma) (15) long after the injury has taken place.

## Results

### Ocular injury leads to infiltration and permanent engraftment of blood-derived monocytes into the retina

We recently showed that corneal alkali injury results in peripheral monocyte infiltration to the retina within 24 hours (1). These cell adopt a dendritiform morphology within 7 days and appear to integrate into the retina (1). In order to determine the long-term effect of the peripheral monocyte infiltration into the retina we employed a CX3CR1^+/EGFP^::CCR2^+/RFP^ bone marrow chimera model, that fate maps CCR2^+^ monocyte derived CX3CR1^+^ cells. Myelodepletion was performed with busulfan, instead of gamma irradiation, since busulfan treatment was recently shown not to cause peripheral monocyte infiltration into the retina of bone marrow transferred mice.(1) To minimize confounding, the contralateral eye of each mouse was used as internal control.

In the absence of retinal injury peripheral CX3CR1^+^ cells do not physiologically infiltrate into the normal retina 16 months after bone marrow transfer (BMT) (fig. 1 A). Only a small number of CX3CR1^+^ cells is present around the optic nerve head (**red arrows)** and along the retinal vessels (**white arrows**), (fig. 1 A, E). However, peripheral CCR2^+^ monocytes are present in the inner retina (fig. 1 B, C, F), appearing as ‘patrolling’ retinal monocytes, corroborating our earlier findings (1). In contrast, after ocular injury, the retina is populated almost entirely (~93%) by peripheral CX3CR1^+^ cells (fig. 1 E), that appear to permanently (16 months post injury) engraft into the tissue (fig. 1 G, H). Peripheral CX3CR1^+^ cells were highly dendritiform, morphologically resembling microglia. These cells migrated and occupied all three distinct retinal microglia strata [ganglion cell layer (GCL), inner nuclear layer (INL), and outer plexiform layer (OPL)] in an orderly fashion, as microglial cells do (fig. 1 I-K), (**Supplemental Video 1**). The extent of peripheral monocyte presence in the retina, 16 months after the injury, appears to causes permanent retinal neuroglia remodeling that does not involve increase in CCR2^+^ cell number (fig. 1 F, L, M). Peripheral CCR2^+^ cells were shown to occupy the three distinct retinal microglia strata (ganglion cell, inner nuclear, and outer plexiform layers) (fig. 1 N-P) without expressing CX3CR1 marker (fig. 1 Q, R). These cells represent a population of peripheral CCR2^+^ monocytes that can mature to CCR2^-^CX3CR1^+^ tissue resident macrophages in the setting of ocular injury (1). These cells appear to resume an apparent ‘patrolling’ function long after removal of the noxious stimuli.

**Figure 1.**
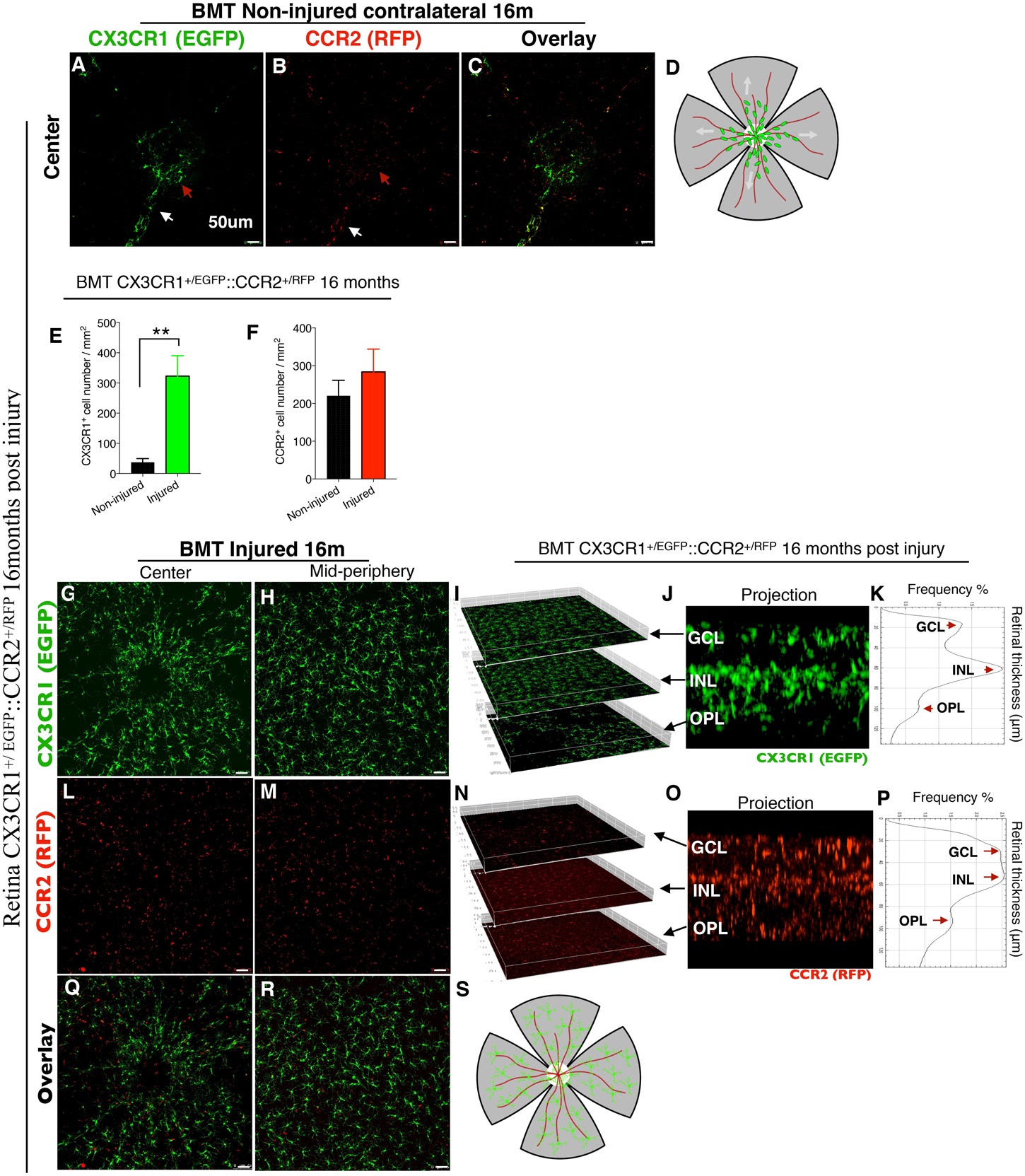
Ocular injury causes permanent engraftment of peripheral CX3CR1^+^ cells in the retina which adopt microglia morphology. Confocal microscopy of busulfan myelodepleted and bone marrow transferred mice with CX3CR1^+/EGFP^::CCR2^+/RFP^ cells 16 months after corneal alkali burn injury. **(A)** Maximum projection of confocal images shows that CX3CR1^+^ cells cannot enter into the retina under physiological conditions. Few cells are present around the optic nerve head (red arrow) and along the retinal vessels (white arrow). **(B)** Maximum projection of confocal images shows CCR2^+^ cells scattered across the retina, located mainly in the GCL. **(C)** Overlay image of CX3CR1 and CCR2 channels. **(D, S)** Representative schematic of flat mount retina as used for confocal imaging; center white circle represents the optic nerve head, red lines the retinal vessels, and green dots the peripheral monocytes. **(E, F)** Quantification of the number of peripheral CX3CR1^+^ and CCR2^+^ cells 16 months after ocular injury shows marked increase in CX3CR1^+^ cells (P<0.01) but normal numbers of CCR2^+^ cells. **(G, H)** Ocular injury causes infiltration and permanent engraftment of peripheral CX3CR1^+^ cells in the retina. **(I)** Isometric reconstruction of confocal stacks separated by retinal layers (GCL, INL, and OPL). **(J)** Cross section of the retina shows presence of CX3CR1+ cells in the CGL, INL, and OPL. **(K)** Histogram of CX3CR1^+/EGFP^ expression within the retinal tissue shows intensity peaks of EGFP signal in the GCL, INL, and OPL. **(I-K)** Sixteen months after the injury, infiltrated CX3CR1^+^ cells migrate into the three distinct microglia stratums (GCL, INL, OPL) and adopt microglia morphology (dendritiform). **(L, M)** Patrolling CCR2^+^ cells are present in the retina but do not differentiate to CX3CR1+ cells. **(N-P)** Even though the number of peripheral CCR2^+^ cells is is normal, peripheral CCR2^+^ cells migrate into the three distinct microglia stratums (GCL, INL, OPL) 16 months after the injury. **(Q, R)** Overlay image of CX3CR1 and CCR2 channels 16 months after ocular injury. GCL: ganglion cell layer, INL: inner nuclear layer, OPL: outer plexiform layer, BMT: bone marrow transfer. **(A-C, G, H, J, M, Q, R)** Scale bar: 50μm. n=5 per group. **(E, F)** Independent t-test. *\*\* P<0.01*.

### Infiltrated peripheral CX3CR1^+^ cells remain pro-inflammatory months after engraftment despite their ramified quiescent morphology

We previously showed that peripheral monocyte infiltration into the retina is associated with marked neuroretinal tissue damage.(1) Prompt inhibition of the monocyte infiltration, using the tumor necrosis factor alpha (TNF-α) inhibitor infliximab, leads to neuroretinal protection (1). In order to assess the nature of engrafted peripheral CX3CR1^+^ monocyte into the retina, we employed a BMT model and flow cytometry to assess MHC-II expression, as an indicator for activation. Five months after acute injury, 72% of the CX3CR1^+^ CD45^hi^ were MHC-II^hi^, despite their otherwise quiescent morphology (fig. 2 A, B). At the same time, 79% of the CX3CR1^+^ CD45^lo^ had no MHC-II expression (fig. 2 A, B). Engrafted peripheral CX3CR1^+^ cells despite their otherwise ramified and quiescent morphology, were shown to ensheath β3-tubulin^+^, suggesting that they are involved in active phagocytosis of of the neuroretinal tissue (fig. 2 C-E) and potentially contributing to neuroretinal degeneration.

**Figure 2.**
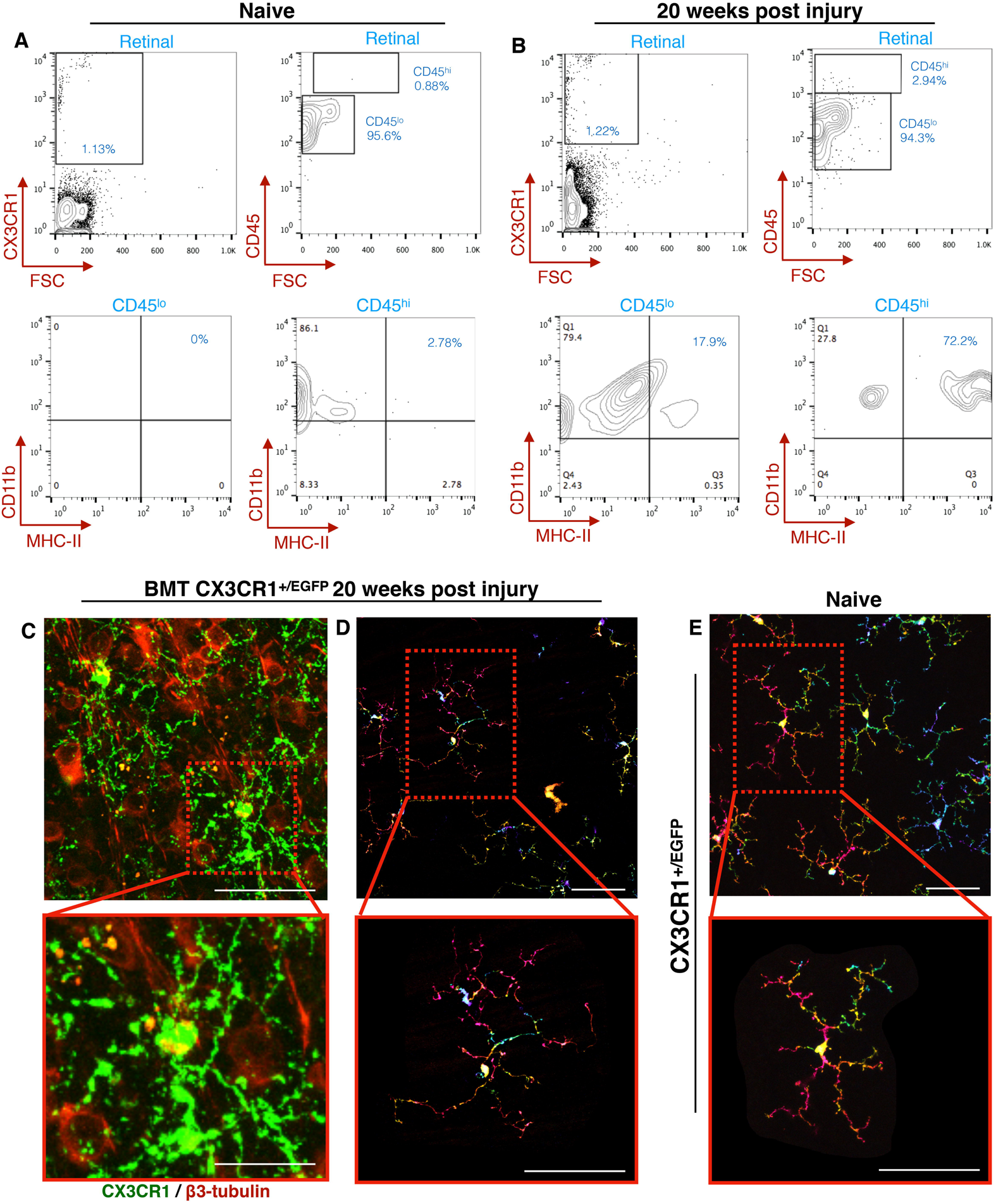
Engrafted peripheral CX3CR1^+^ cells remain reactive despite their morphometric quiescence. Flow cytometric analysis of retinal cells from CX3CR1^+/EGFP^ mice. **(A)** In naive mice, CX3CR1^+^ cells are predominantly CD45^lo^ (96%) and only a small percentage is CD45^hi^ (4%). CX3CR1^+^ CD45^lo^ cells do not express MHC-II, however, 2.8% of CX3CR1^+^ CD45^hi^ have baseline MHC-II expression. **(B)** Twenty weeks after ocular injury, 94% percent of the CX3CR1^+^ are CD45^lo^ of which 20% express MHC-II. Likewise, 3% of the CX3CR1^+^ are CD45^hi^ of which 72% express MHC-II. **(C)** Twenty weeks after the injury, peripheral CX3CR1^+^ cells appear to frequently interact and ensheath β-III tubulin^+^ neuroretinal tissue, **(C-E)** despite their otherwise ramified morphology. Scale bars: **(C-D)** top panel=50μm, **(C-D)** bottom panel= 20μm, **(D, E)**. All experiments were triplicates.

### Ocular injury causes peripherally engrafted microglia to become resistant to CSFR1 inhibition

To characterize further the nature of the retina engrafted monocytes after ocular injury, we performed microglia ablation using systemic administration of PLX5622, a small molecule that inhibits colony stimulating factor 1 receptor (CSF1R) and leads to microglia depletion.(16, 17). First, we assessed the ability of PLX5622 to deplete retinal microglia. Administration of PLX5622 for 3 weeks, completely removed retinal microglia from CX3CR1^+/EGFP^ mice. This event was followed by a gradual repopulation of the retina within 3 weeks. The repopulating cells migrated into the three distinct retinal microglia strata (GCL, INL, OPL) and transformed to a dendritiform morphology. PLX5622 treatment was performed in bone marrow transfer mice to assess peripheral monocyte contribution in microglia repopulation (fig. 3 A). These repopulating cells were primarily from the periphery (fig. 3 B-N) with a small number (~7%) originating from non-peripheral microglia progenitors, whose contribution increased after acute ocular injury (~20%) (**Supplemental Figure S1 A-G**). Repopulating cells migrates in the three distinct microglia strata within 6 weeks of PLX treatment (fig. 3 O-Q). These repopulated cells remained sensitive to CSF1R inhibitor and could be re-ablated by re-administration of PLX5622 (fig. 3 R-V). In contrast, in the setting of ocular injury, peripherally engrafted monocytes were resistant to CSF1R inhibitor and could not be ablated with PLX5622 (Fig 4 A-H). These cells appeared semi-ramified, occupying all three distinct microglia strata (fig. 4 I-K), but with higher density in the OPL (fig. 4 K) likely do to local migration. The resistance of retinal microglia that populated the retina after ocular injury was not attributed to their peripheral origin, since repopulated microglia in the absence of ocular injury also originated from the periphery (fig. 3 F, L) and remained sensitive to PLX5622 inhibition after engraftment (fig. 3 S, T, V).

**Figure 3.**
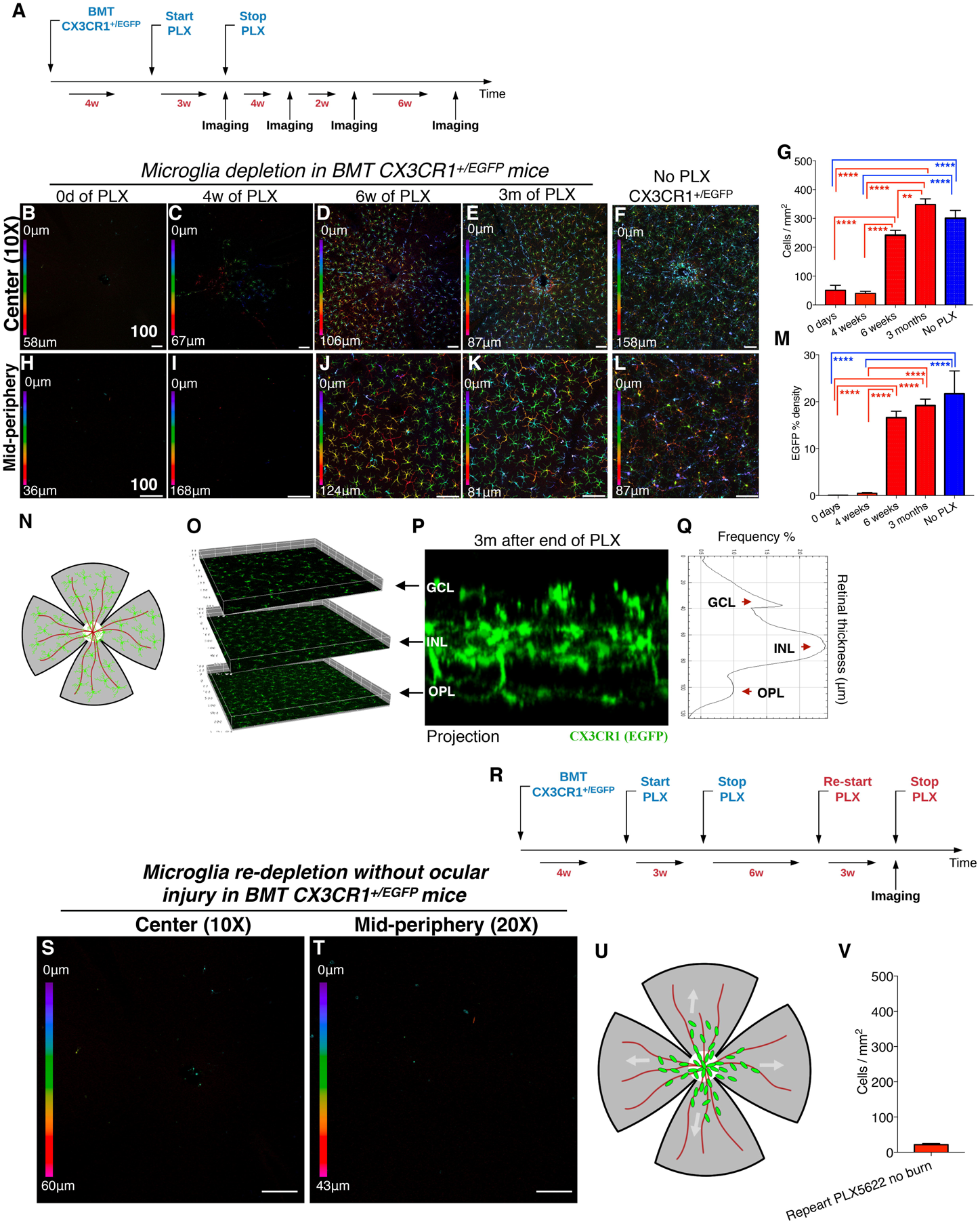
Microglia repopulation after PLX5622 treatment occurs primarily by peripheral CX3CR1^+^ cells. **(A)** Busulfan myelodepleted C57BL/6 mice received adoptive transfer of CX3CR1^+/EGFP^ bone marrow cells. Four weeks later they received PLX5622 treatment for 3 weeks and followed by confocal microscopy 3 weeks after cease of PLX5622 treatment. **(B-E)** Central color-depth coded confocal images of flat mount retinas 0, 3, 7, and 21 days after cease of PLX5622 treatment and **(F)** of naive CX3CR1^+/EGFP^ mice. **(G)** Quantification of CX3CR1^+^ cell number. **(H-K)** Mid-peripheral color-depth coded confocal images of flat mount retinas 0, 3, 7, and 21 days after cease of PLX5622 treatment and **(L)** of naive CX3CR1^+/EGFP^ mice. **(M)** Quantification of EGFP intensity. **(B, F, G, H, M)** PLX5622 treatment for 3 weeks does not cause acute peripheral CX3CR1^+^ cell infiltration into the retina. **(D, G, J, M)** However, 6 weeks after cease of PLX5622 treatment peripheral CX3CR1^+^ cells appeared to occupy the majority of the retina area, **(E, G, K, M)** and at 3 months, the retina is completely repopulated by peripheral CX3CR1^+^ cells. **(G, M)** The number of repopulated peripheral CX3CR1^+^ cells at 3 months after the end of PLX5622 treatment approximates that of naive CX3CR1^+/EGFP^ mice with no PLX5622 treatment. **(N, U)** Representative schematic of flat mount retina as used for confocal imaging; center white circle represents the optic nerve head, red lines the retinal vessels, and green dots the peripheral monocytes. **(O)** Repopulating peripheral CX3CR1^+^ cells migrated into all three distinct microglia strata (GCL, INL, OPL) 3 months after the injury. **(P)** Cross section of the retina shows presence of CX3CR1+ cells in the CGL, INL, and OPL. **(Q)** Histogram of CX3CR1^+/EGFP^ expression within the retinal tissue shows intensity peaks of EGFP signal in the GCL, INL, and OPL. **(R)** Re-application of PLX5622 in repopulated peripheral microglia. **(S, T)** Peripheral monocytes the repopulate the retina remain sensitive to CSF1R inhibition and can be re-depleted by re-application of PLX5622 treatment. (V) Quantification of peripheral CX3CR1^+^ cells after re-application of PLX5622 shows complete depletion of repopulated CX3CR1^+^ cells. n=5 per group. GCL: ganglion cell layer, INL: inner nuclear layer, OPL: outer plexiform layer, BMT: bone marrow transfer. **(B-I, J, L)** Scale bar: 100μm. Multiple comparisons using Tukey’s method \*\**P<0.01*, \*\*\*\**P<0.0001*.

**Figure 4.**
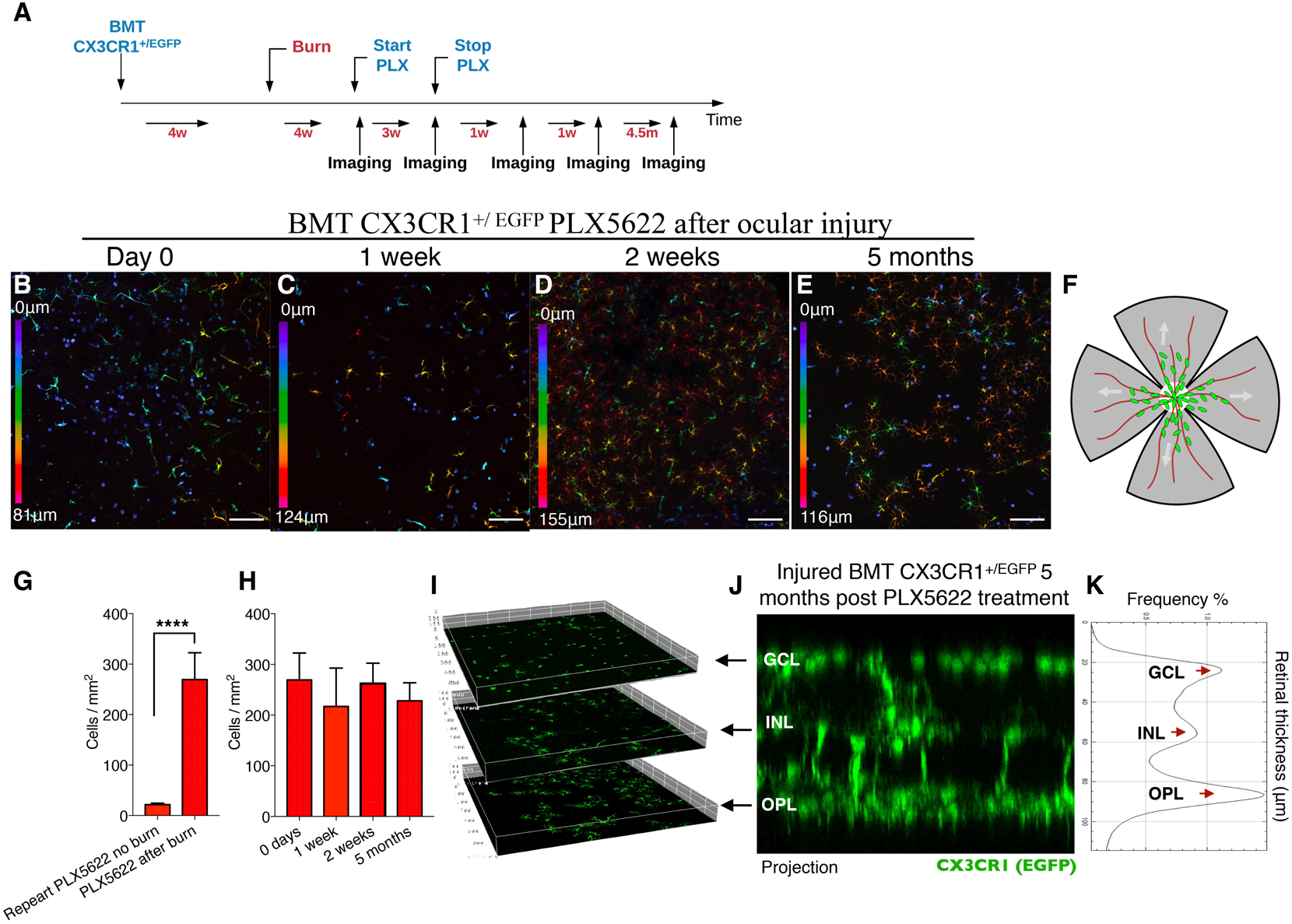
Ocular injury leads to population of the retina by peripheral CX3CR1^+^ that are resistant to CSF1R inhibitor. **(A)** Busulfan myelodepleted CX3CR1^+/EGFP^ bone marrow chimeras received ocular injury and 4 weeks later PLX5622 treatment for 3 weeks. **(B)** Immediately after cease of PLX5622 treatment peripherally populated CX3CR1^+^ cells remained present in the retina and **(G)** resistant to CSF1R inhibition, as pose to peripherally repopulated CX3CR1^+^ cells in the absence of ocular injury. **(F)** Representative schematic of flat mount retina as used for confocal imaging; center white circle represents the optic nerve head, red lines the retinal vessels, and green dots the peripheral monocytes.**(B)** Inhibition resistant CX3CR1^+^ cells were localized primarily in the GCL, with fewer cells in the INL and OPL. **(C)** 1 week, **(D)** 2 weeks, and **(E)** 5 months after cease of PLX5622 treatment peripherally populated CX3CR1^+^ cells remained present in the retina **(H)** with unchanged cell number. **(I, J)** However, over the 5 months peripheral CX3CR1^+^ cells have migrated into all 3 distinct microglia strata, **(K)** with unusually increased cell density in the OPL. GCL: ganglion cell layer, INL: inner nuclear layer, OPL: outer plexiform layer, BMT: bone marrow transfer. n=3 per group. **(B-E)** Scale bar: 100μm. **(G)** Independent t-test. **** *P<0.0001*.

## Discussion

Here, we have shown that ocular injury not only causes peripheral monocyte infiltration into the retina, as previously demonstrated (1), but leads to permanent engraftment and remodeling of the neuroglia system to a pro-inflammatory state resistant to CSFR1 inhibition. This process is characterized by a) infiltration of peripheral CX3CR1^+^CCR2^+^ cell into the retina within hours after the injury, b) migration into the three distinct microglia strata (GCL, INL, OPL), c) differentiation to dendritiform cells, and finally d) permeant engraftment into the retina as tissue resident CX3CR1^+^CCR2^−^ monocytes. Interestingly, despite their integration into the tissue and morphometric resemblance to yolk-sac derived microglia, these cells remain different from their predecessors. Contrary to native microglia, peripherally engrafted monocytes retain expression of activation marker MHC-II, are actively involved in phagocytosis of the neuroretinal tissue, and become resistant to CSF1R inhibitor despite their otherwise quiescent morphology and integration into the retina.

Using a small molecule inhibitor (PLX5622) of CSF1R (an essential receptor for microglia survival) we show that engrafted peripheral CX3CR1^+^ monocytes after injury differ from the yolk-sac derived microglia, and cannot be ablated using CSF1R inhibition. In contrast, peripheral CX3CR1^+^ cells repopulating in the absence of ocular injury remain sensitive to PLX5622 treatment and can be re-ablated. On one hand, the findings could be attributed to the progeny of the infiltrating cell-type and perhaps the existence of bone marrow cells that give rise to post-embryonic self-maintaining tissue resident macrophages. On the other hand, it could be due to epigenetic effects stemming from the injury leading to changes in gene expression and subsequent changes in phenotype. Indeed, epigenetic reprogramming can occur in brain microglial after exposure to inflammatory mediators, such as TNF-α, IL-1β, and IL-6 (18). This effect persist for months and leads to microglia immune training and tolerance. Therefore, altering microglia epigenetic reprogramming through cytokine inhibition may be a potential therapeutic avenue for retina and brain neuropathology (15, 18).

Previous studies have shown that infiltrated peripheral monocytes in the CNS remain transcriptionally similar to peripheral myeloid cells 7 days after injury (19) and distinctly different to microglia (5, 20-24). The lack of adaptation of recruited peripheral monocytes in the CNS environment may contribute to functional neuroglia changes and subsequent damage to the tissue (19). Whether CSF1R resistance is attributed to epigenetic reprogramming or selective recruitment of distinct progenitor cells that do not express CSF1R is not known and merits further investigation. However, recent reports have demonstrated the existence of two waves of temporally separated and functionally distinct monocyte progenitors that emerge from the yolk-sac between E7.5 and E.8.5 (25). The first wave consists of CSF1R^hi^-c-Myb^−^ monocyte progenitors give rise mostly to local macrophages and microglia without monocyte intermediates. The second wave consists of CSF1R^lo^c-Myb^+^ monocyte progenitors which give rise locally to yolk-sac macrophages. Most of these latter cells migrate into the fetal liver through the blood circulation at E9.5 to generate cells of multiple lineages (25). They likely represent cells that transiently become progenitors and precursors for the majority of tissue-resident macrophages through a monocytic intermediate (25, 26) which could give rise to tissues resident macrophages post-embryonically (25). From our study, it appears that local cues, such as chemokine and cytokines expression in the retina (1), may be pivotal in determining the type of EMP cell (early or late) recruited into the adult retina, without excluding the possibility that both EMP waves originate from a single population that matures into CSF1R^+^ and CSF1R^−^ cells (25).

Regardless of the mechanism of cell selection or differentiation of EMPs, the presence of peripheral monocytes in the retina constitutes a major homeostatic change (1, 8, 12, 13). Infiltration of bone-marrow derived myeloid cells was previously described in disease models such as amyotrophic lateral sclerosis(27), Alzheimer disease (28), scrapie (29), bacterial meningitis (30), and other diseases of the CNS (4). Peripheral monocytes have also been shown to contribute to nerve axon degeneration 42 days after spinal cord injury (21). Our recent work demonstrated that MHC-II^+^ peripheral monocytes infiltrate into the retina promptly after corneal injury, and these cells mediate inflammation and subsequent neuroretinal tissue damage (1). Here we show that peripheral monocytes that infiltrate into the retina remain MHC-II positive months after infiltration, and despite their otherwise quiescent morphology, engage in phagocytosis of neuroretinal tissue. Quiescent microglia have low MHC-II expression, while activated microglia and infiltrated macrophages express high MHC-II (1, 8, 31-34). Prolonged MHC-II expression by retinal microglia contributes to neuropathology in multiple sclerosis (35), Alzheimer’s (36), and Parkinson’s disease (37). Furthermore, gradual increase in MHC-II expression in the white matter has been shown to cause myeline loss and cognitive decline in aged human and nonhuman primates (38). Our findings are in agreement with previous studies showing that the presence of peripheral monocytes in the CNS and retina is pathologic and likely contributes to neuroinflammation (1, 8, 12, 13). Prolonged residency of activated monocytes in the retina can lead to progressive gradual neuronal damage and irreversible visions loss, which recapitulates the clinical picture of glaucomatous damage after anterior segment injury, and may appear clinically as loss of retina ganglion cells. Under normal intraocular pressure this can resemble glaucoma seen in patients with chemical injuries (39-41), and could explain the increased glaucoma susceptibility in patients that have suffered acute ocular injuries (2, 3, 14, 15). In the setting of ocular injury, ramified CX3CR1^+^ cells from the bone marrow may occupy and engraft into the three distinct microglia strata, thereby co-existing with embryonically derived microglia, yet as we have shown in this study they have distinct pro-inflammatory phenotype. Thus, modifying these immunological processes, either by inhibiting the infiltration of peripheral monocytes or re-training them to suppress their inflammatory phenotype may be have therapeutic implications for patients suffering for neurodegenerative diseases secondary to inflammation and injury.

Another important finding of this study is the involvement of microglia in the regulation of peripheral monocyte infiltration into the retina. Microglia depletion, by using PLX5622, initiates a ‘stress signals’ that causes retina microglia repopulation from peripheral CX3CR1^+^ monocytes that are CSF1R^+^. Conversely, in the absence of PLX5622 treatment or ocular injury, peripheral CX3CR1^+^ cells are unable to infiltrate into the retina, and their presence is limited to around the optic nerve head (1, 8). It is important to mention that microglia depletion with PLX5622 is a non-physiological event which disturbs homeostasis, and, therefore, likely contributes to the deregulation of the neuroglia system. Indeed, microglia depletion with PLX5622 leads to repopulation primarily from peripheral monocytes (~93%) and secondarily from non-peripheral progenitors (7%). In the setting of ocular injury, the contribution from non-peripheral progenitors increases to 20%, still relatively low. The inability of non-peripheral microglia progenitors to supply the required cells for prompt repopulation may be attributed to the nature of microglia, which are long lived, self-renewing cells (24), and therefore have a low turn-over rate (42). In the setting of acute microglia depletion, peripheral monocytes seem to be necessary to support the process of repopulation and overcome the slow turn-over rate of microglia progenitors. The ability of CSF1R independent embryonic precursors to functionally replace yolk sac derived microglia during development has been demonstrated (26), suggesting that other tissue macrophages may assume the role of microglia under certain conditions. Our findings are in disagreement with a recent report which showed that peripheral monocytes have little contribution to microglia repopulation following PLX5622 treatment (43). Both studies confirm the existence of two distinct CX3CR1^+^ cells populations (peripheral and central) but their relative contribution differs. Perhaps differences in the fate mapping techniques used may explain this disparity. We used busulfan myelodepletion which was previously shown not to result in artificial peripheral monocyte infiltration into the retina (1, 4) whereas the contrasting study (43), employed a non-BMT CX3CR1^creERT2^-tdTomato model in which most peripheral tdTomato^+^ monocytes are lost 3 moths after tamoxifen induction due to their high turn-over rate (44) while microglia retain tdTomato due to their low turn-over (42, 45). However, it is possible that peripheral monocytes may arise from a specific peripheral monocyte sub-population that has not lost tdTomato expression and live significantly longer once engrafted into the tissue. Further studies are needed to resolve this discrepancy.

In conclusion, we have demonstrated that a surface injury to the eye can cause not only peripheral monocyte infiltration into the retina, but also permanent engraftment and integration into the neuroretinal tissue; morphologically resembling yolk sac-derive microglia. However, despite this resemblance, peripherally derived microglia are functionally distinct from yolk sac derived ones, remaining long-term activated expressing MHC-II, involved in active phagocytosis of neuronal tissue, and becoming resistant to CSF1R-inhibition. These features are not attribute to their peripheral origin, in the absence of ocular injury, peripherally monocytes that repopulate the retina after PLX5622 treatment, remain sensitive to CSF1R inhibition and MHC-II^lo^. This observed microglia remodeling contributes to progressive neurodegeneration, which clinically recapitulates the glaucomatous damage to the retina observed in patients with similar ocular injuries. Therefore, modifying this phenotype of peripheral monocytes that contribute to inflammatory microglial remodeling may have clinical therapeutic implications in diseases where continued neuroinflammatory processes contribute to neurodegeneration.

## Materials and Methods

### Mouse model of alkali burn

All animal-based procedures were performed in accordance with the Association For Research in Vision and Ophthalmology (ARVO) Statement for the Use of Animals in Ophthalmic and Vision Research, and the National Institutes of Health (NIH) Guidance for the Care and Use of Laboratory Animals. This study was approved by the Animal Care Committee of the Massachusetts Eye and Ear Infirmary. Mice were bred in house (Massachusetts Eye and Ear Animal Facility) and used at the ages of 6-12 weeks. CX3CR1^CreERT2^::ROSA26-tdTomato mice were generated by breeding male B6.129P2(C)-Cx3cr1^tm2.1(cre/ERT2)Jung^/J with female B6.Cg-Gt(ROSA)26Sor^tm14(CAG-tdTomato)Hze^/J. CX3CR1+/E^GFP/^::CCR2^+/RFP^ were generated by breeding male B6.129(Cg)-*Ccr2^tm2.1lfc^*/J with female B6.129P-*Cx3cr1^tm1Litt^*/J. The breeding schemes are described in **Supplemental Figure 2**. The following strains were obtained from Jackson Laboratory (Bar Harbor, ME, USA): C57BL/6J (Stock# 000664) B6.129P-*Cx3cr1^tm1Litt^*/J (CX3CR1^EGFP/EGFP^) (Stock# 005582),(46) B6.129(Cg)-*Ccr2^tm2.1lfc^/J (CCR2*^RFP/RFP^) (Stock# 017586),(47) *B6.129P2(C)-CX3CR1^tm2.1(cre/ ERT2)Jung^/J* (Stock# 020940(48) and B6.Cg-Gt(ROSA)26Sor*^tm14(CAG-tdTomato)Hze^/J* (Stock# 007914) (49).

Alkali chemical burns were performed according to our previous study (14). In brief, mice were anesthetized using ketamine (60 mg/kg) and xylazine (6 mg/kg) and deep anesthesia was confirmed by toe pinch. Proparacaine hydrocloride USP 0.5% (Bausch and Lomb, Tampa, FL, USA) eye drop was applied to the cornea, and after 1 minute was carefully dried with a Weck-Cel (Beaver Visitec International, Inc, Waltham, MA, USA). A 2 mm diameter filter paper was soaked in 1 M sodium hydroxide (NaOH) solution for 10 seconds, dried of excess alkali and applied onto the mouse cornea for 20 seconds. After the filter paper was removed, prompt irrigation with sterile saline for 10 seconds was applied. The mouse was then placed on a heating pad, positioned laterally, and the eye irrigated for another 15 minutes at low pressure using sterile saline. Buprenorphine hydrochloride (0.05 mg/kg) (Buprenex Injectable, Reckitt Benckiser Healthcare Ltd, United Kingdom) was administered subcutaneously for pain management. A single drop of topical Polytrim antibiotic was administered after the irrigation (polymyxin B/trimethoprim, Bausch & Lomb Inc, Bridgewater, NJ, USA). Mice were kept on the heating pad until fully awake.

### Tissue preparation for flat mount imaging

Following the alkali burn, eyes were enucleated at predetermined time points and fixed in 4% paraformaldehyde (PFA) solution (Sigma-Aldrich, St Louis, MO, USA) for 1 hour at 4°C. The cornea and retina tissue were carefully excised using microsurgical techniques and washed 3 times for 5 minutes in phosphate buffer solution (PBS) (Sigma-Aldrich, St Louis, MO, USA) at 4°C. The tissues were then incubated in 5% bovine serum albumin (Sigma-Aldrich, St Louis, MO) and permeabilized using 0.3% Triton-X (Sigma-Aldrich, St Louis, MO, USA) for 1 hour at 4°C. The specific antibody was added in blocking solution, incubated overnight at 4°C, and then washed 3 times for 10 minutes with 0.1% Triton-X in PBS. The tissues then were transferred from the tube to positive charged glass slides using a wide pipette tip with the concave face upwards. Four relaxing incisions from the center to the periphery were made to generate 4 flat tissue quadrants. VECTRASHIELD^®^ mounting medium (Vector Laboratories, H-1000, Vector Laboratories, CA, USA) was placed over the tissue followed by a coverslip.

### Retinal flat mount imaging

The tissues were prepared for flat mount and imaged using a confocal microscope (Leica SP-5, Leica Microsystems inc, Buffalo Grove, IL, USA). Images were taken at 10x, 20x, 40x and 63x objective lenses using z-stack of 0.7, 0.6, 0.4 and 0.3 μm step, respectively. Image processing with ImageJ version 1.5s was used to obtain maximum and average projections of the z-axis, color depth maps, and 3-D volumetric images. The number of retinal microglia/macrophage cells and their density was quantified using imageJ software.

### Bone marrow chimera model

C57BL/6J mice were myelodepleted with 3 intraperitoneal injections of busulfan, an alkylating agent that depletes bone marrow cells (Sigma-Aldrich, St Louis, MO, USA) (35mg/kg) 7, 5, and 3 days prior to bone marrow transfer. CX3CR1^+/EGFP^::CCR2^+/RFP^, B6::CX3CR1^+/EGFP^ and CX3CR1^CreERT2^::ROSA26-tdTomato bone marrow cells (5×10^6^ total bone marrow cells) were transferred to the myelodepleted C57BL/6J mice through tail vein injection 1 month prior to corneal alkali burn. Bactrim (trimethoprim/sulfamethoxazole resuspended in 400mL drinking water) was given *ad lib* for 15 days post busulfan treatment.

### Flow cytometric analysis of retinal cells

Eyes were harvested, retinas were isolated surgically, digested with Papain Dissociation System (Worthington Biochemical Corporation, Lakewood, NJ, USA) and processed with flow cytometry. Retinal microglia/macrophage cells were detected as EGFP^+^ or tdTomato population in CX3CR1^+/EGFP^ and CX3CR1^CreERT2^::ROSA26-tdTomato mice, respectively. Different fluorescent CD45 (Clone: 104), CD11b (Clone: M1/70), I-A/I-E (MHC-II) (Clone: M5/114.15.2), CX3CR1 (Clone: SA011F11), Ly6C (Clone: HK1.4) and Ly6G (Clone: 1A8) antibodies (Biolegend, San Diego, CA, USA) were used to perform the flow cytometry. Cells were analyzed on a BD LSR II cytometer (BD Biosciences, San Jose, CA, USA) using FlowJo software (Tree Star, Ashland, OR, USA).

### Microglia depletion

Microglia depletion was perform using PLX5622, a CSF1R inhibitor. The compound was provided by Plexxikon Inc. (Berkeley, CA, USA) and formulated in AIN-76A standard chow by Research Diets Inc. (New Brunswick, NJ, USA). A dose of 1200 ppm was given to mice for 3 weeks prior to analysis.

### Statistical analysis

Results were analyzed with the statistical package of social sciences (SPSS) Version 17.0 (Statistical Package for the Social Sciences Inc., Chicago, IL, USA). The normality of continuous variables was assessed using the Kolmogorov-Smirnov test. Quantitative variables were expressed as mean ± standard error of mean (SEM) and qualitative variables were expressed as frequencies and percentages. The Mann-Whitney test was used to assess differences between the groups. All tests were two-tailed, and statistical significance was determined at p < 0.05. The independent student t-test was used to compare between two groups, and pairwise t-test to compare changes within the same group. Analysis of variance (ANOVA) was used for comparison of multiple groups. Alpha level correction was applied, as appropriate, for multiple comparisons.

## Author contributions

EIP designed the research study, conducted experiments, acquired data, analyzed data and wrote the manuscript. FL, CZ, and VK conducted experiments, acquired data, analyzed data and reviewed the manuscript; DV, RD, JC and CHD performed critical review of the manuscript.

## Acknowledgments

This work was supported by the Boston Keratoprosthesis Research Fund, Massachusetts Eye and Ear, the Eleanor and Miles Shore Fund, the Massachusetts Lions Eye Research Fund, an unrestricted grant to the Department of Ophthalmology, Harvard Medical School, from Research to Prevent Blindness, NY, NY, and NIH National Eye Institute core grant P30EY003790.

## Figures and Legends

**Supplemental Figure S1. Microglia repopulation after PLX5622 treatment occurs primarily by peripheral CX3CR1^+^ cells.**

**(A-C)** Flat mount retinas from busulfan myelodepleted *CX3CR1^CreERT2^::ROSA26-tdTomato* bone marrow transfer mice were stained with Iba1-FITC 2 months after PLX5622 treatment. Cre-Lox recombination was induced with 5 daily tamoxifen injections, 1 weeks prior to analysis. Two months after PLX5622 treatment, retinal microglia repopulation occurs by peripheral tdTomat^+^ FITC^+^ and non-peripheral tdTomat^-^FITC^+^ CX3CR1^+^ cells. (D) Flow cytometry analysis of retinal cells 2 months after PLX5622 treatment confirms that microglia repopulation occurs primarily by peripheral CX3CR1^+^ cells and at a smaller percentage (7%) by non-peripheral CX3CR1^+^. **(E)** Representative schematic of flat mount retina as used for confocal imaging; center white circle represents the optic nerve head, red lines the retinal vessels, and green dots the peripheral monocytes. **(F)** In the setting of ocular injury, the percentage of non-peripheral CX3CR1^+^ cells that repopulate the retina increases to 20%. n=3 per group. **(A-C)** Scale bar: 100μm.

**Supplemental Figure S2. Mouse breeding schemes.**

**(A)** CX3CR1^+/EGFP^ **(B)** CX3CR1^+/EGFP^::CCR2^+/RFP^, and **(C)** CX3CR1^CRE/ERT2^::R26tdTomato reporter mice were were generated by in house using the depicted breeding schemes. **(D)** Explanation of mouse strain nomenclature, as provided by Jackson Laboratory.

**Supplemental Video 1.** Peripheral CX3CR1^+^ cells engraft permanently in the retina after corneal injury.

CX3CR1^+/EGFP b^ one marrow chimera model 16 months after transfer. In control eye, few CX3CR1^+/EGFP^ cells from the blood enter into the retina through the retinal vessels and the optic nerve head. These cells are primarily localized in the ganglion cell layer, and do not migrate deeper into the tissue (left video). Corneal alkali burn causes vast infiltration of blood-derived CX3CR1^+/EGFP^ monocytes that permanently engraft in the retina. Sixteen months after bone marrow transfer, infiltrated CX3CR1^+/EGFP^ cells are highly ameboid and migrate into the three distinct retinal microglia stratums (right video). The morphology and topology of the infiltrated peripheral CX3CR1^+/EGFP^ cells resembles that of the retinal microglia.

